# AMBIENT: Accelerated Convolutional Neural Network Architecture Search for Regulatory Genomics

**DOI:** 10.1101/2021.02.25.432960

**Authors:** Zijun Zhang, Evan M. Cofer, Olga G. Troyanskaya

## Abstract

Convolutional neural networks (CNN) have become a standard approach for modeling genomic sequences. CNNs can be effectively built by Neural Architecture Search (NAS) by trading computing power for accurate neural architectures. Yet, the consumption of immense computing power is a major practical, financial, and environmental issue for deep learning. Here, we present a novel NAS framework, AMBIENT, that generates highly accurate CNN architectures for biological sequences of diverse functions, while substantially reducing the computing cost of conventional NAS.

## 1 Introduction

Convolutional neural networks (CNNs), which led to breakthroughs in computer vision research as part of the 2012 ImageNet competition[1] and the ensuant rise of deep learning[2], have found extensive use in the domain of regulatory genomics[3, 4]. A significant amount of work in this area has focused on predicting the presence of regulatory features (e.g. transcription factors, histone marks, chromatin accessibility) directly from genomic sequence inputs[5–10].

As manually designing the network architecture is a time-consuming process with significant implications for model performance, we recently developed AMBER[11] to automate the process by extending computer vision neural architecture search (NAS) algorithms[12, 13] to apply them to the regulatory feature prediction domain. In essence, AMBER enables the trading of computing power for accurate CNN architectures without large time commitments or a deep understanding of machine learning.

However, the computing power required for deep learning, and for NAS in particular, can cause time delays and incur high financial and environmental costs[14]. For instance, it is estimated that training a state-of-the-art natural language processing deep learning model with NAS emits as much carbon dioxide as five cars over their entire life-time[14]. Thus, there is an urgent need for more emission-efficient and environmentally-friendly NAS methods. Furthermore, despite intense work on the regulatory feature prediction problem, it is unclear whether different regulatory features should be modeled with different CNN architectures.

We herein describe a new method, AMBIENT, which addresses this challenge by incorporating information about the regulatory features in the search process. For a given regulatory feature’s dataset (e.g. ChIP-seq peaks), AMBIENT maps a summary of that dataset to the initial state of the controller model and generates an optimal task-specific architecture. We show that AMBIENT is more efficient than existing methods, allowing it to identify architectures of comparable accuracy at an accelerated pace.

## 2 Methods

### 2.1 Overview of AMBIENT framework

Generally, NAS methods consist of a controller model and its child models [12]. Given a task of interest, the controller model generates child neural architectures from a model search space, and the sampled architectures are subsequently implemented, trained, and evaluated for their prediction performance. The feedback loop between the controller-generated architectures and their performance evaluations facilitates the training of the controller model to generate better child architectures. Existing NAS approaches[11–13] designate a one-to-one correspondence between a controller and a child task during the controller training.

In this work, we augment the controller model to generate multiple task-specific architectures based on the distinct characteristics of said tasks, as illustrated in Fig. 1. The introduction of data descriptors enables the controller to jointly learn the optimal architectures among diverse and similar tasks of interest. During application, AMBIENT generates an optimal architecture for a new task based on its data descriptors, instead of training from scratch, thus substantially reducing its need for computing power. Mathematical and technical details for the AMBIENT controller model are in Appendix A.

**Figure 1:**
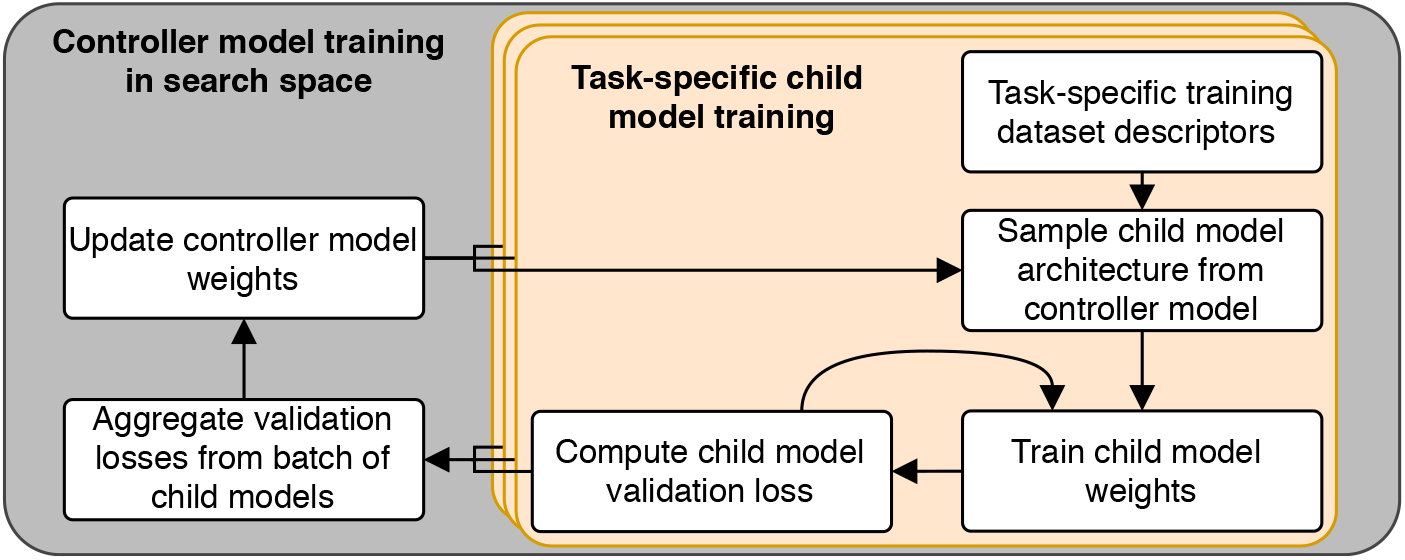
Visual overview of the AMBIENT method.

### 2.2 Compilation of Dataset and Data Descriptors

Training and testing data were taken from the original publication introducing CNNs for the regulatory feature prediction task[6]. From the original set of 919 regulatory features, we randomly drew 36 for training. As this work is still in the pilot stages, we set aside the remaining data for more extensive analysis at a later date. Regulatory features were held out while still taking into consideration bias due to cell-type specific coverage similarity (i.e. chromatin accessibility and RNA polymerase features were held-out on a per-tissue basis).

To summarize each regulatory feature, we included a number of data descriptors. First, we considered high-level data descriptors such as whether the regulatory feature in question was a transcription factor, RNA polymerase, histone mark, or chromatin accessibility. Based on the InterPro domains[15] present in the training set of regulatory features, we added two data descriptors indicating whether a given regulatory feature possessed a C2H2-type zinc-finger domain or a C2H2 bZIP Maf domain. In addition to these coarse-grained data descriptors, we also considered a number of more fine-grained quantitative measures. We included descriptors for the number of peaks in a given sequencing experiment, as well as the median and variance of peak widths. To gauge the locality of each regulatory feature, we recorded the distance from each peak to the nearest annotation for several annotations – splice sites, exons, genes, and UTRs – from GENCODE v34[16]. These distances were summarized with their median and variance. We also included the average conservation across the regulatory regions using phastCons[17] and phyloP[18]. Finally, we included quantitative summaries of motifs identified in each regulatory feature. In specific, we used HOMER 2[19] to identify motifs of 6, 8, and 10 bp, and then measured the multiplicity and strand bias for each motif length considered. We summarized both measures with their minimum, maximum, and median. For the multiplicity, we also included the variance.

### 2.3 Design of the Model Search Space

We designed a 10-layer model search space to systematically evaluate the optimal architecture differences among distinct epigenetic regulatory features. Specifically, the 10 layers were divided into three consecutive convolution-pooling-dropout blocks, where higher blocks simultaneously increased the channel dimension while reducing the feature dimension by a factor of 4, a common heuristic used in computer vision. The number of channels in the blocks were set to 16, 64, and 256, respectively. Within each block, there were seven candidate convolution operations with hyperparameters of kernel size *k* ∈ {1, 8, 14, 20} and dilation rate^1^ *d* ∈ {1, 2}. For the pooling portion of the block, there were two feature-level pooling operations: max-pooling and average-pooling. Both pooling operations used a step size of 4 and stride of 4. Finally, each block had a dropout operation, with dropout rates *p* ∈ {0.1, 0.3, 0.5}. In the last block, we replace the feature-level pooling with four distinct channel-level pooling operations: flatten, global max or average-pooling, or a LSTM.

## 3 Results

### 3.1 Training AMBIENT on Diverse Epigenetic Features Reveals Both Distinct and Universal Aspects of Optimal Neural Architectures

We trained AMBIENT jointly with 36 epigenetic regulatory features. For each individual regulatory feature, we sampled neural architectures from the trained controller RNN using its corresponding data descriptors as inputs, then grouped the architectures based on their biological functions (i.e. transcription factor, RNA polymerase, histone mark, or chromatin). Strikingly, our marginalized analysis demonstrates that AMBIENT learned distinct neural architectures for modeling genomic sequences regulated by different epigenetic features (Fig. 2). For example, to model genomic sequences with DNase activity, AMBIENT strongly prefers average pooling after convolution with large kernel size (k=20) as the first two layers; by contrast, sequences bound by TFs are considered to be better modeled with max pooling following convolution with large kernel size. Similarly, sequences bound by RNA polymerase exclusively prefer a smaller kernel size for the first convolution operation, supported by previous work[5]. Within each functional category, we observed relatively smaller variations of architectures for particular factors (data not shown).

**Figure 2:**
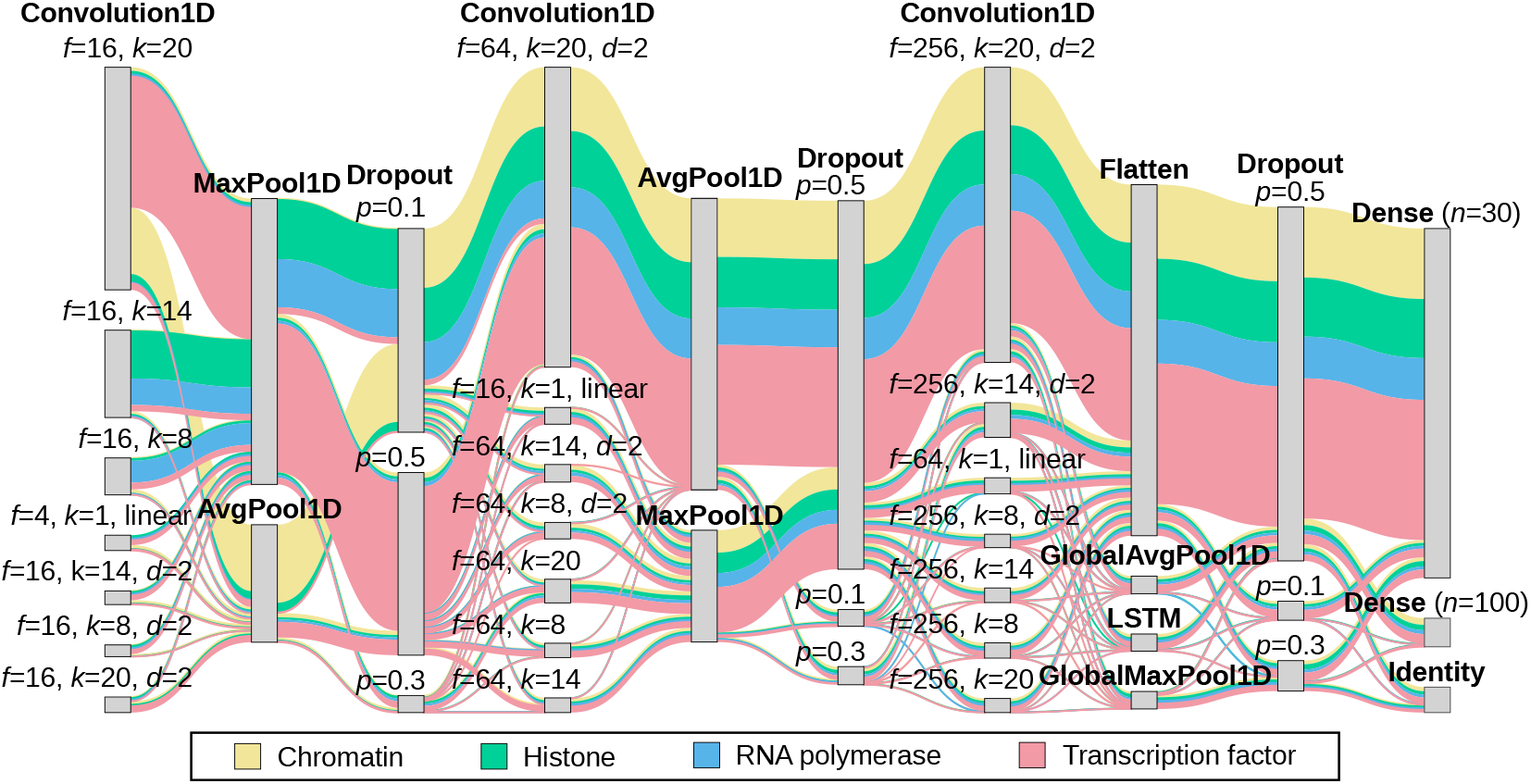
Sankey diagram of the results for different classes of regulatory features. Convolutional and dense layers are followed with ReLU activation layers unless otherwise noted (e.g. linear). *n* is the number of output units for a dense layer. For a dropout layer, *p* is the dropout rate. For a convolutional layer, *f* is the number of filters, *k* is the kernel width, and *d* is the dilation. Note that *d* = 1 for convolutional layers unless stated otherwise.

Interestingly, the variable architectures are predominantly in the first convolution-pool-dropout block, whereas the latter two blocks are largely constant across regulatory features, indicating partial neural architectures, especially top layers after initial feature extraction, are transferable across vastly diverse genomic sequence contexts.

### 3.2 AMBIENT Predicts Accurate Neural Architectures with Minimal Computing Power Overhead

A big challenge for NAS is its requirement of high computing power, and more recently uncovered, its huge emission footprint [14]. Thus, we sought to evaluate the quality of AMBIENT-generated neural architectures on held-out regulatory features, and how much runtime and computing power is reduced without training NAS from scratch. To that end, we ran the single-run AMBER genomics NAS method[11] on four held-out epigenetic features, one for each functional category (Fig. 3). We analyzed the average performance of architectures generated by AMBER over time. We directly compared these performance measurements with the performance of models generated by the trained AMBIENT controller model (i.e. applying AMBIENT directly to the held-out dataset descriptors without any fine-tuning). Indeed, comparable performances on the validation datasets were achieved for both AMBER and AMBIENT, indicating both methods optimized the neural architectures and were upper-bounded inherently by the training dataset. However, AMBIENT only required a single forward-pass, and thus had minimal computing power overhead. In comparison, AMBER had to be trained from 8 to 21 GPU hours, with an average of 15 GPU hours, to achieve comparable or better performance.

**Figure 3:**
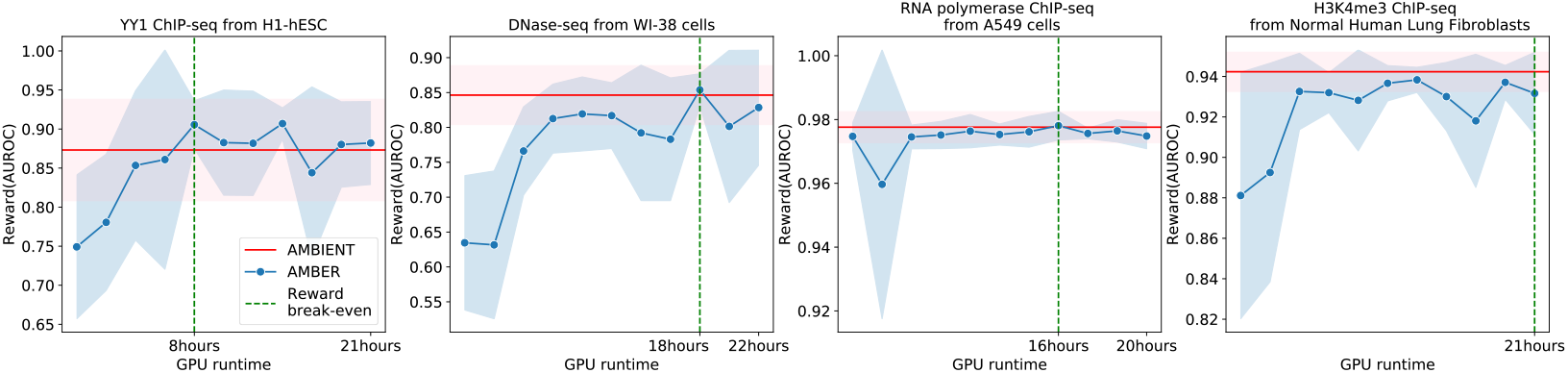
Comparison of test performance and GPU runtime on a single NVIDIA V100 GPU. For each functional category, one representative held-out independent regulatory genomics dataset was analyzed by running AMBER[11] (blue), and applying the trained AMBIENT directly, to infer the neural architecture using the held-out dataset descriptors (red). Lines are average rewards for *n* = 10 architectures generated by AMBER or AMBIENT, trained independently; shaded areas are 95% confidence intervals; reward break-even points are the first time that AMBER rewards exceed that of AMBIENT.

## 4 Conclusions

We present a novel NAS method, AMBIENT, that can generate accurate CNN architectures capable of modeling genomic sequences for diverse epigenetic regulatory features, while substantially reducing the carbon-emission footprint. Analysis of AMBIENT-designed neural architectures can provide valuable hypotheses for testing in computational and functional genomics.

## Acknowledgments and Disclosure of Funding

The authors thank all members of the Troyanskaya lab for their helpful discussions. The authors are also pleased to acknowledge that this work was performed using the high-performance computing resources at the Simons Foundation and the TIGRESS computer center at Princeton University. E.M.C. was supported by the National Science Foundation Graduate Research Fellowship Program (NSF-GRFP) and National Institutes of Health (NIH) grant T32 HG003284. This work was supported by NIH grant R01 GM071966 (O.G.T.). O.G.T. is a senior fellow of the Canadian Institute for Advanced Research (CIFAR) Genetic Networks program.

## Appendix A AMBIENT Controller Recurrent Neural Network

For any given child neural network, we can model its network architecture as a sequence of computational layers that maps input features to output labels, where the operation per layer is determined in an auto-regressive manner. Following the existing work[11–13], we employ a Recurrent Neural Network (RNN) to capture the layer-wise dependencies:

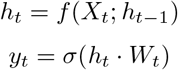

We parameterize *f* with a long short-term memory network (LSTM); for each layer *t*, its computational operations are sampled from a multinomial distribution *y_t_*, specified by a softmax transformation *σ*(·) of *h_t_* · *W_t_*. Given *y_t_*, we sample the categorically-encoded architecture token *X*_*t*+1_ and feed it to update the LSTM hidden states for step *t* + 1.

Unlike conventional NAS optimizations, where the initial LSTM state *h*_0_ is random initialized and subsequently learned through optimization, we aim to incorporate the dataset-specific descriptors to inform the initial state: *h*_0_ = *g*(*D_k_*), where *D_k_* is a vector of data descriptors for the *k*-th dataset, and *g* is parameterized as a 16-unit multi-layer perceptron that maps *D_k_* to *h*_0_.

The controller parameters *θ* are learned by maximizing the validation AUC, denoted by *R_k_*, across a set of *K* datasets with distinct data descriptors {*D_k_*|*k* = 1, 2, .., *K*}. Let the likelihood of selecting a target architecture *X* given the parameters *θ* be denoted as *π*(*X*; *θ*): *π*(*X*; *θ*) = ∏_*t*_ ℙ(*X_t_*|*X*_(*t*−1):1_; *θ*)).

Since *R_k_* is not differentiable with respect to the controller parameters *π*(*X*; *θ*), we employ a Reinforcement learning framework to maximize validation AUC as a reward. In this work, we obtain the policy gradient by minimizing the surrogate loss function by Proximal Policy Optimization (PPO) [20]. Specifically, using a batch of architectures and their reward signals, the loss function is:

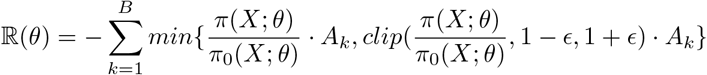

where *ϵ* is the clipping strength and set to 0.2 following the previous report[20]; *A_k_* is the advantage for the *k*-th target architecture. We estimate *A_k_* by subtracting the corresponding moving average of the *k*-th dataset’s reward, denoted by *b_k_*; followed by re-scaling to have the unit variance within a batch: 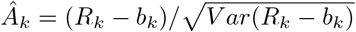.

The advantage estimates 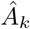 are inserted in ℝ(*θ*) to obtain the policy gradients. In practice, the estimation for *A_k_* is crucial for training the AMBIENT controller RNN, because it needs to normalize the reward signals over multiple datasets with potentially different validation AUC baselines and variances.

To compare the efficiency of AMBIENT, we also run the plain version of AMBER[11] without the consideration of data descriptors, which we refer to as single-run AMBER because it runs on a single regulatory epigenetic feature. For fair comparison, we did not employ the parameter-sharing module when asking the AMBER modeler to implement a child architecture[13]. Optimization-related parameters (e.g. LSTM units, learning rate, batch size) for the controller RNN were set to be identical between AMBIENT and single-run AMBER. We trained AMBIENT using 4 Nvidia-v100 GPUs and single-run AMBER using a single Nvidia-v100 GPU.

## Footnotes

1 Note that dilation rate is not applicable when *k* = 1

